# Spectrum of pathogenic variants and multiple founder effects in amelogenesis imperfecta associated with *MMP20*

**DOI:** 10.1101/2020.07.23.217927

**Authors:** Georgios Nikolopoulos, Claire E. L. Smith, James A. Poulter, Gina Murillo, Sandra Silva, Catriona J. Brown, Peter F. Day, Francesca Soldani, Suhaila Al-Bahlani, Sarah A. Harris, Mary J. O’Connell, Chris F. Inglehearn, Alan J. Mighell

## Abstract

Amelogenesis imperfecta (AI) describes a heterogeneous group of developmental enamel defects that typically have Mendelian inheritance. Exome sequencing of ten families with recessive hypomaturation AI revealed 4 novel and 1 known variants in the matrix metallopeptidase 20 (*MMP20*) gene that were predicted to be pathogenic. *MMP20* encodes a protease that cleaves the developing extracellular enamel matrix and is necessary for normal enamel crystal growth during amelogenesis. New homozygous missense changes were shared between four families of Pakistani heritage (c.625G>C; p.(E209Q)) and two of Omani origin (c.710C>A; p.(S237Y)). In two families of UK origin and one from Costa Rica, affected individuals were homozygous for the previously reported c.954-2A>T; p.(I319Ffs*19) variant. For each of these variants, microsatellite haplotypes appeared to exclude a recent founder effect, but elements of haplotype were conserved, suggesting more distant founding ancestors. New compound heterozygous changes were identified in one family of European heritage; c.809_811+12delACGgtaagattattainsCCAG; p.(?) and c.1122A>C; p.(Q374H). All four new variants are within the zinc dependant peptidase domain. This report further elucidates the mutation spectrum of *MMP20* and the probable impact on protein function, confirms a consistent hypomaturation phenotype and shows that mutations in *MMP20* are a common cause of autosomal recessive AI in some communities.

**Data Availability:** The data that support the findings of this study are openly available in ClinVar at https://www.ncbi.nlm.nih.gov/clinvar/, accession numbers: SCV001338799 - SCV001338802 and in the AI Leiden Open Variation Database (LOVD) at http://dna2.leeds.ac.uk/LOVD/ with reference numbers: 0000000313 – 0000000317.

## Introduction

The hardness of dental enamel is remarkable and is a result of its high mineral and low protein content (C. E. Smith, 1998). The process of enamel formation, amelogenesis, begins with production of a secreted soft enamel protein matrix which is then transformed into a mature, organized structure of rod and interrod enamel crystallites almost devoid of protein. Crucial to this transformation process is the secretory stage enamel protease, matrix metallopeptidase 20 (MMP20) [NM_004771] (MIM #604629).

MMP20 is a zinc dependent endopeptidase that is secreted in trace amounts during the secretory and transition stages of amelogenesis by ameloblasts (Llano et al., 1997; Seymen et al., 2015). Once activated, it selectively cleaves the secreted enamel proteins amelogenin, enamelin and ameloblastin, into several products with distinct functional roles (Simmer and Hu, 2002). MMP20 is also thought to facilitate ameloblast movement during secretion, through cell-cell communication, and may influence ameloblast gene expression (Guan and Bartlett, 2013). These activities create newly voided space within which enamel crystallites are able to grow in width and thickness, and finally, to interlock (Bartlett, 2013). If they are not removed, the enamel proteins occupy the enamel volume and restrict the growth of crystallites, leading to brittle enamel that fails prematurely.

Mutations in *MMP20* and nineteen other genes have been shown to cause non-syndromic AI with autosomal dominant, autosomal recessive or X-linked inheritance (J. Kim et al., 2019; C. E. L. Smith et al., 2017; C. E. L. Smith et al., 2020 and http://dna2.leeds.ac.uk/LOVD/genes), while perhaps as many again have been implicated in syndromic AI (Dubail et al., 2018; J. T. Wright et al., 2015). AI, therefore, describes a heterogeneous group of conditions characterized by inherited enamel defects in both dentitions. It can present as a hypoplastic phenotype, where a deficit in secretion results in the formation of a reduced volume of mineralized enamel, or as hypomineralised AI, whereby failure in maturation results in enamel of full thickness but which is soft or brittle and fails prematurely (Gadhia et al., 2012). Hypomineralised AI can be further sub-classified as hypomaturation (brittle enamel), caused by incomplete removal of protein from the developing enamel, or hypocalcification (soft enamel), caused by insufficient transport of calcium ions into the forming enamel (C. E. L. Smith et al., 2017), though these phenotypes often overlap. Mutations in *MMP20* cause autosomal recessive hypomaturation AI. 14 pathogenic variants have been reported to date (Supp. Table 1), most of which are located in the catalytic domain of MMP20 and are thought to affect the stability and functionality of the protein structure.

This study describes the identification of pathogenic *MMP20* variants in ten families segregating autosomal recessive hypomaturation AI, providing additional insights into the spectrum of *MMP20* mutations and associated phenotype. Four novel variants are reported, two of which are relatively common founder mutations in specific populations. Additionally, we perform molecular dynamic simulations of variants in the catalytic domain of MMP20, to determine the effect that these changes have on the protein. This study increases the total reported pathogenic *MMP20* variants to seventeen, suggests that defects in *MMP20* are a more common cause of AI than was previously reported and enriches our understanding of the effects that mutations in the catalytic domain of MMP20 can have on its functionality.

## Materials & Methods

### Patients

Individuals from each of the ten families were recruited following informed consent in accordance with the principles outlined in the declaration of Helsinki, with local ethical approval (REC 13/YH/0028). A diagnosis of AI was made by a dentist after clinical examination, based on the physical appearance of the dentition, and was confirmed by dental X-ray. Genomic DNA was obtained via venous blood samples, using conventional extraction techniques, or from saliva using Oragene^®^ DNA Sample Collection kits (DNA Genotek, ONT, Canada) and extracted according to the manufacturer’s instructions.

### Whole-exome sequencing and analysis

Three micrograms of genomic DNA from a single individual from each family (marked with an arrow on the pedigrees) was subjected to whole exome sequencing (WES) and analysed as described previously (C. E. L. Smith et al., 2019). Variants present in the dbSNP150 database of NCBI or the Genome Aggregation Database (gnomAD; http://gnomad.broadinstitute.org) with a minor allele frequency (MAF) ≥1% were excluded. The potential pathogenic effect of the variants was predicted using Combined Annotation Dependent Depletion (CADD v1.3, https://cadd.gs.washington.edu) (Rentzsch et al., 2019), the Sorting Intolerant From Tolerant algorithm (SIFT, Sim et al., 2012), PROtein Variation Effect ANalyzer (PROVEAN, Choi et al., 2012), and MutPred2 (http://mutpred.mutdb.org/index.html). The potential effect on splicing for each intronic variant was predicted by the Human Splicing Finder v3.1 (http://umd.be/HSF3/HSF.shtml).

### PCR and Sanger Sequencing

Mutations were confirmed and segregation performed for all available family members, marked with (*) on each pedigree of Figure 1. Primer sequences can be found in Supp. Table 2. Sanger sequencing was performed using the BigDye Terminator v3.1 kit (Life Technologies, CA, USA) according to manufacturer’s instructions and resolved on an ABI3130xl sequencer (Life Technologies). Results were analysed using SeqScape v2.5 (Life Technologies).

**Figure 1.**
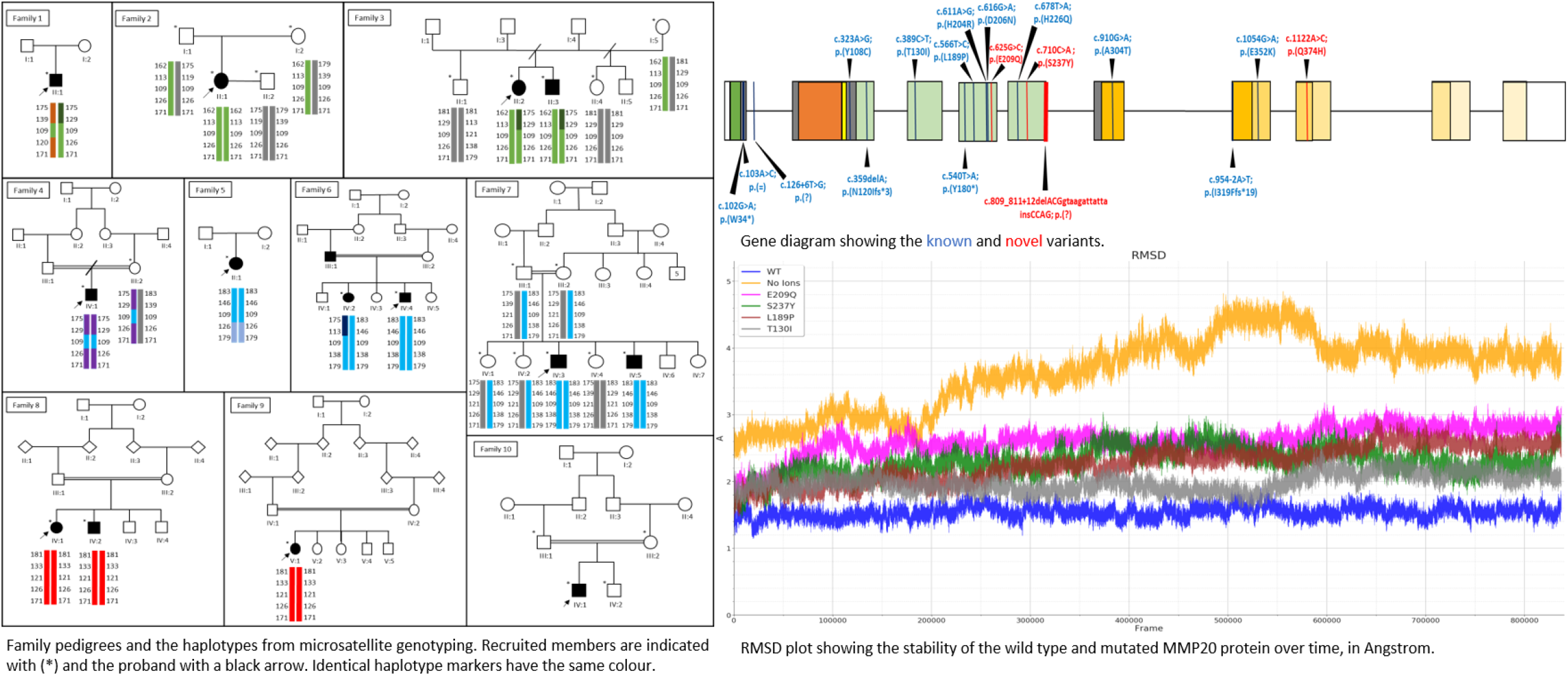
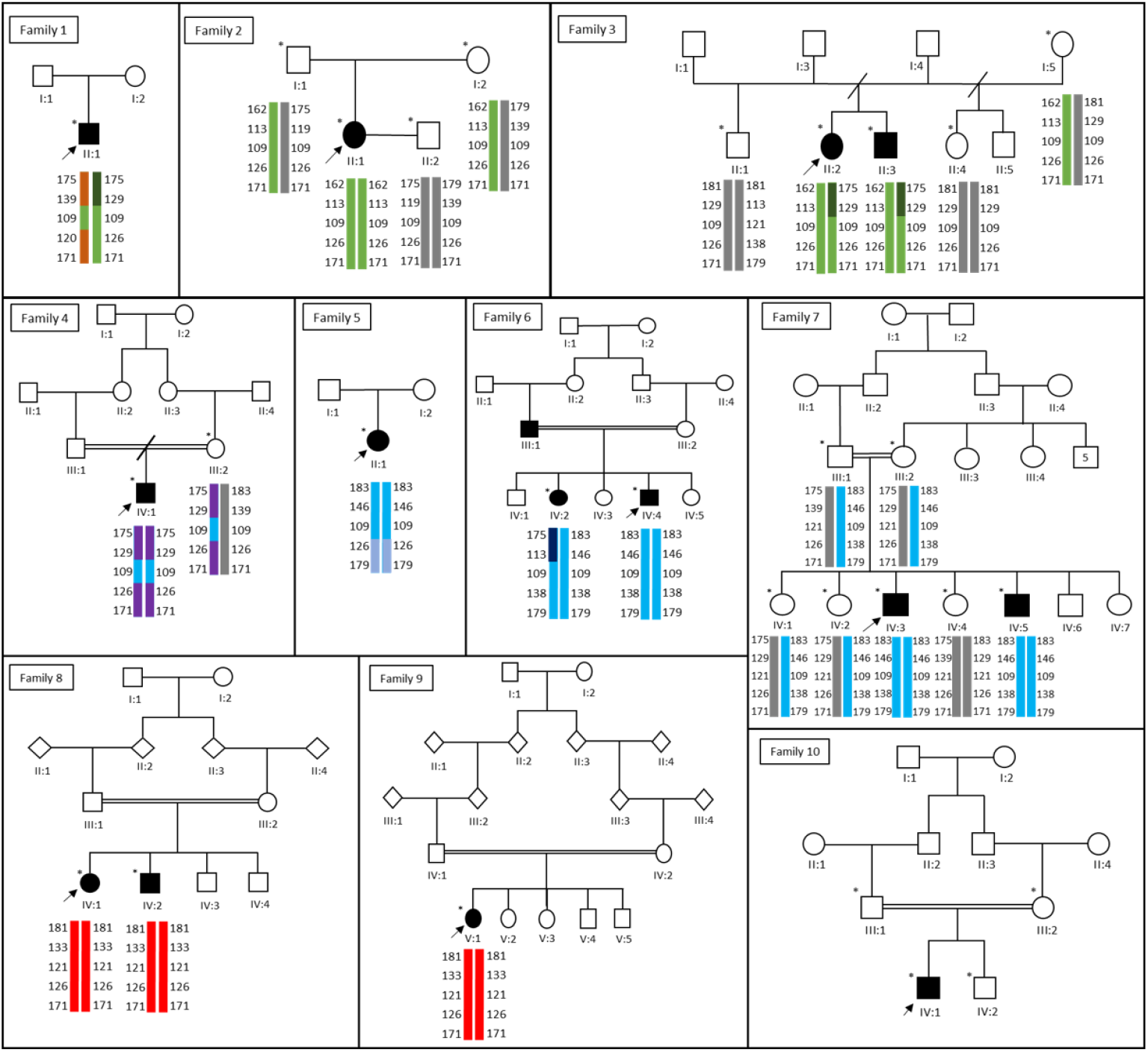
Pedigrees of the ten families recruited for this study. The genotypes determined from the microsatellite markers are presented beneath each individual examined. Families 1, 2 and 3 carry the c.954-2A>T variant, Families 4, 5, 6 and 7 carry the c.625G>C variant, Families 8 and 9 carry the c.710C>A variant and Family 10 has the c.809_811+12delACGgtaagattattainsCCAG and c.1122A>C variants. The common haplotypes are presented in the same colour and the haplotypes without a pathogenic variant are coloured grey. The markers are presented in the order: 11cen, D11S940, D11S1339, *MMP20*, D11S4108, D11S4159, D11S4161, 11qter. The recruited family members are marked with an asterisk and the proband in each family is indicated with a black arrow.

### Microsatellite analysis

Primer sequences for markers were obtained from the UCSC genome browser and standard HEX-tagged primers (Sigma-Aldrich, Supplementary Table 2) were used to assess the flanking haplotypes of *MMP20*. The analysis was performed as described previously (Nikolopoulos et al., 2020).

### Protein Structure Analysis

The tertiary structure of MMP20 has been determined in atomistic detail by nuclear magnetic resonance (NMR) (PDB: 2JSD) (Arendt et al., 2007), which provides the starting structure for molecular dynamics (MD) simulations. Amino acid substitutions were made in the Wild Type (WT) structure using the Chimera visualization tool (Pettersen et al., 2004). MD simulations were performed using AMBER18 (Case et al., 2018). Protocols used to perform the MD simulations are in the Supplementary Methods. Rhapsody was used for *in silico* saturation mutagenesis analysis (Ponzoni et al., 2020) and the prediction of the changes in solvent accessibility, residue occlusion and free energy (ΔΔG) of the mutated proteins was performed with Site Directed Mutator (SDM) (Pandurangan et al., 2017).

## Results

Ten unrelated families presenting with features consistent with autosomal recessive hypomaturation AI, in the absence of any co-segregating disease, were recruited to the study (Figures 1 & 2). Genomic DNA from affected members of each family was subjected to exome sequencing. Detailed coverage statistics can be found in Supp. Table 3. Variant files were filtered to select rare variants with high predicted pathogenicity, then variants in known AI causing genes were highlighted. This revealed biallelic mutations in *MMP20* (Supp. Table 4) in each family. The position of these variants in the gene is shown in Figure 3, along with representative electropherograms for each novel variant. PCR and Sanger sequencing confirmed the variants segregated with AI in all available family members.

**Figure 2.**
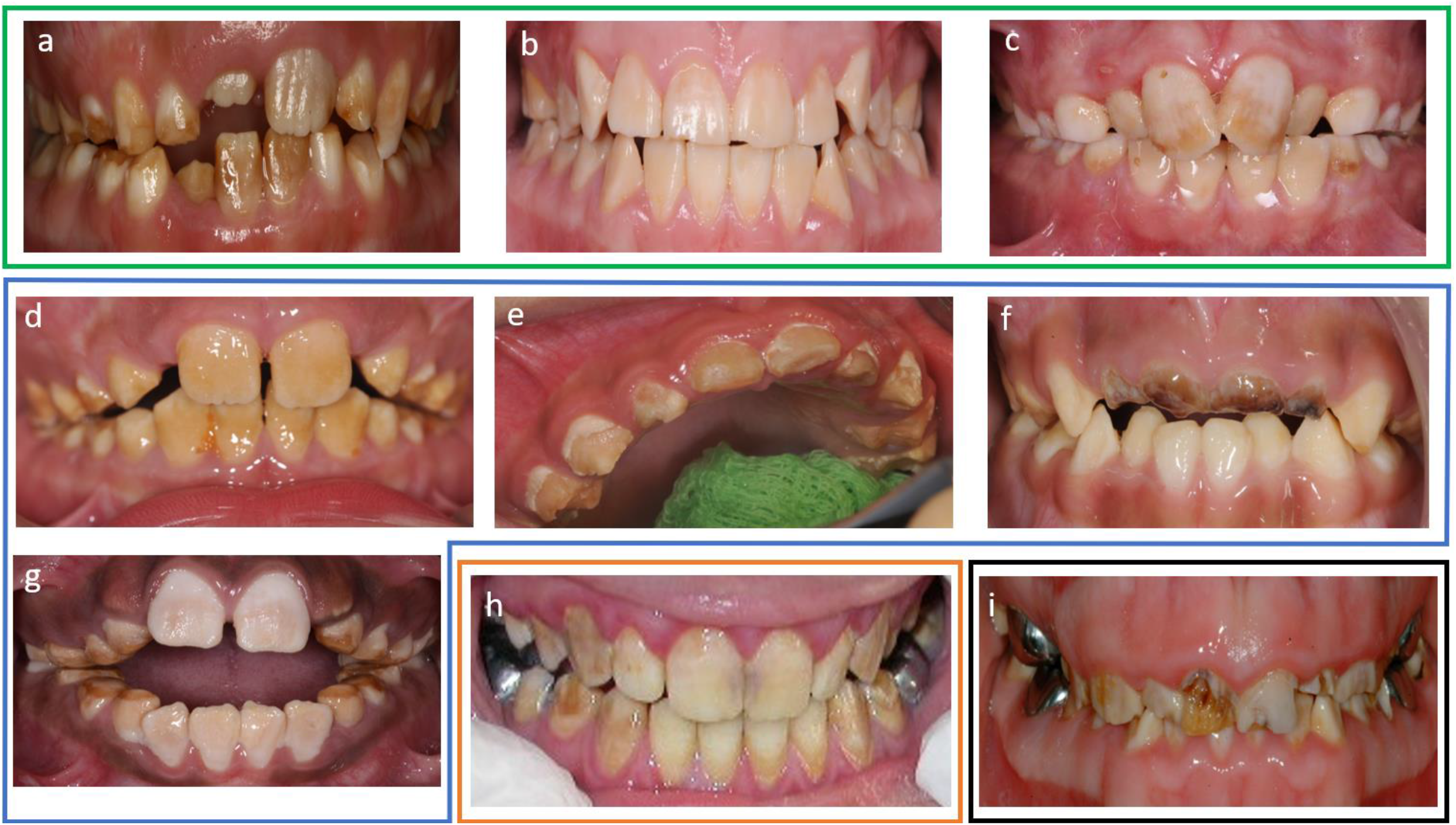
Clinical images for nine probands at different chronological ages, illustrating the spectrum of variations that can be observed within the overall, typical features of hypomaturation AI. Panels a–c are Families 1–3, who all have European heritage and share the same homozygous *MMP20* c.954-2A>T; p.(I319Ffs*19) change. Panels d–g are Families 4–7, who all have Pakistani heritage and share the same homozygous *MMP20* c.625G>C; p.(E209Q) change. Panel h is Family 9 who along with Family 8 (no clinical images available) have Omani heritage and have the same homozygous *MMP20* c.710C>A; p.(S237Y) homozygous change. Panel i is Family 10, of European heritage, which was characterised by the compound heterozygous *MMP20* c.809_811+12delACGgtaagattattainsCCAG; p.(?) and c.1122A>C; p.(Q374H) changes.

**Figure 3.**
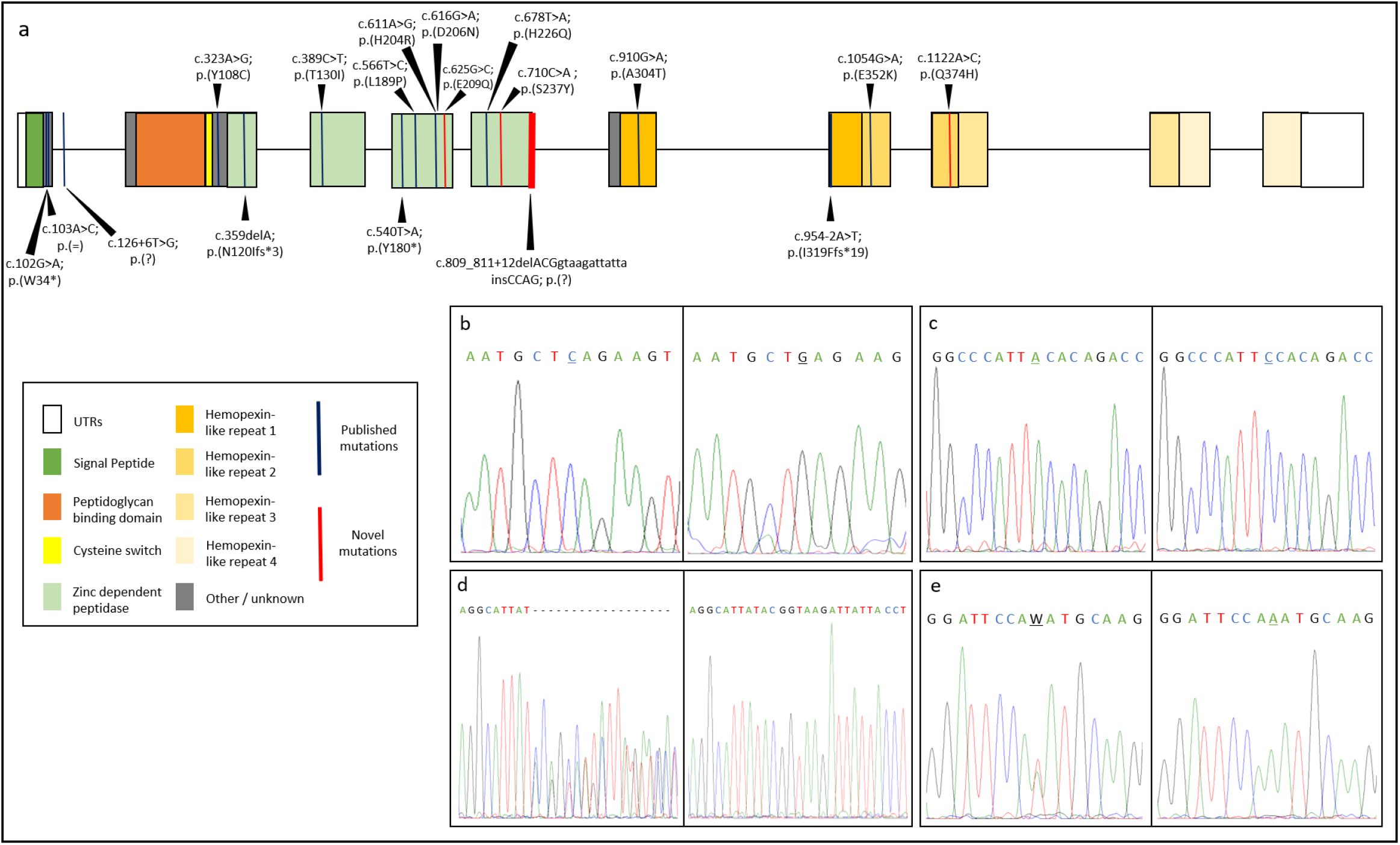
a: *MMP20* gene diagram. The positions of all known mutations are marked on the gene with a vertical bar, novel variants are highlighted in red. The zinc dependent peptidase domain is the catalytic domain of the protein, where most pathogenic variants have been found. b-e: A representative electropherogram of each of the novel variants found in this study, b– c.625G>C variant and WT, c– c.710C>A variant and WT, d– c.810_812+12delACGgtaagattattainsCCAG variant and WT, e– c.1122A>C variant and WT.

In Families 1,2 and 3, a homozygous frameshift variant (NM_004771: c.954-2A>T, NP_004762: p.(I319Ffs*19)) was identified in intron 6. This has been published previously as a cause of autosomal recessive hypomaturation AI and is expected to lead to retention of the sixth intron (J. W. Kim et al., 2005). To confirm this hypothesis, we attempted to perform reverse transcriptase PCR of the *MMP20* transcript on control blood cDNA. However, no amplification was achieved, suggesting that the level of *MMP20* expression in blood is below the threshold for detection.

A novel homozygous missense mutation, c.625G>C, p.(E209Q), was identified in exon 4 as the cause of disease in Families 4, 5, 6 and 7. This is known variant rs199788797, which has not previously been associated with a disease phenotype. In the gnomAD database this variant has a frequency of 0.000457 in the South Asian population but is absent from all other reported populations. E209 is fully conserved in the mammalian clade, as shown in Supp. Figure 1, using the sequences listed in Supp. Table 5. Additionally, the mutation is predicted to be damaging (Supp. Table 6).

In Families 8 and 9, a novel homozygous missense mutation, c.710C>A, p.(S237Y) was identified in exon 5. This variant is absent from databases, is evolutionarily conserved in all the mammalian species analysed (Supp. Figure 1) and is predicted to cause disease by all algorithms used (Supp. Table 6).

The affected individual in Family 10 was found to be a compound heterozygote for a novel missense mutation c.1122A>C, p.(Q374H) in exon 8 and a novel deletion-insertion (delins) variant: c.809_811+12delACGgtaagattattainsCCAG, p(?), spanning the splice donor site of intron 5. Both are absent from variant databases. Variant p.(Q374H) is predicted to be damaging (Supp. Table 6), and Q374 is conserved in the mammalian clade (Supp. Figure 1). Interestingly, mouse *Mmp20* has a histidine in the equivalent position. This presumably does not have a detrimental effect on the function of the mouse protein, perhaps due to other changes of the mouse gene that balance the protein or reflecting differences in enamel development and function between mouse teeth, which grow throughout life, and human teeth which do not. Nevertheless, mutation prediction software does classify this variant as pathogenic, it is absent from the gnomAD database, and paired with a second *MMP20* variant it has given rise to a hypomaturation phenotype consistent with biallelic *MMP20* disease. The delins variant is predicted by Human Splicing Finder to disrupt the intron 5 splice donor site, possibly leading to retention of the fifth intron (Supp. Figure 2).

The families that share variants (Families 1, 2 and 3 with c.954-2A>T, Families 4-7 with c.625G>C and Families 8 and 9 with c.710C>A) also originate from the same ethnic backgrounds. To determine whether these families share common founder haplotypes at the *MMP20* locus, we genotyped five microsatellite markers across a 1.5 cM/3 Mb region of chromosome 11q22 in each family, in the order: 11cen, D11S940, D11S1339, *MMP20*, D11S4108, D11S4159, D11S4161, 11qter. Genotyping results are presented in Supp. Table 7. Haplotype analysis is shown in Figure 1.

In Omani Families 8 and 9, carrying the c.710C>A variant, haplotyping suggests they are closely related. However, the picture is more complex in the remaining two family groups. Families 1-3 (the first from Costa Rica, the remaining two from the UK) carry the previously published c.954-2A>T variant. The proband in Family 2 is homozygous for a haplotype also seen in Family 3, but a second haplotype in Family 3 has a proximal recombination. Family 1 could share the recombinant haplotype observed in family 3 together with an unrelated haplotype, but without phase information this cannot be confirmed. Interestingly, all three families are homozygous for D11S4108 100kb from *MMP20*. Families 4-7, of UK Pakistani origin, carry the c.625G>C variant. The proband in Family 6 is homozygous for the same haplotype segregating in Family 7. However, a second haplotype with a distal recombination is seen in the second affected sibling in Family 6, suggesting the affected (unsampled) father carries both haplotypes. Family 5 is homozygous for a third haplotype, again identical at the proximal end to that in Family 7 but recombinant at the distal end. Family 4 in contrast is homozygous for a fourth haplotype identical to the Family 7 haplotype at the distal end but proximally recombinant. Again, all four families are homozygous for the immediately adjacent marker D11S4108.

The catalytic centre of MMP20 is a 160 residue domain containing the zinc dependent peptidase active site. It contains one catalytic and one structural zinc ion; the structures modelled here additionally contain two calcium ions, which are also structural. We assessed how missense variants affect the protein function by performing MD simulations of the relevant mutated proteins. In addition, to demonstrate the key role of metal ions in maintaining MMP20 structure, we performed MD simulations in which the metal ions were removed. We compared the protein structures and dynamics observed for the WT with metal ions present, with those obtained for p.(E209Q) and p.(S237Y) from this study (see Figure 4a) and p.(T130I) and p.(L189P) from the literature (Figure 4b); and also with results of the WT in the absence of metal ions. Figure 4c shows the root mean squared deviation (RMSD) of each of the simulations from their starting structures (the averages calculated for each repeat are in Supp. Table 8, and the repeats are shown separately in Supp. Figure 3). An increase in RMSD relative to the WT, implying decreased protein stability, was observed for all variants, apart from p.(T130I). We also analysed the changes in three key inter-atomic interactions between the WT and the variants, atomic fluctuations (Supp. Figure 4a), hydrogen bonding interactions (Supp. Table 9) and salt bridges (Supp. Table 10), to provide insight into why these particular variants cause functionally deleterious changes in protein structure. The most significant structural distortions were observed in the simulations performed in the absence of structural zinc and calcium ions (see Supp. Figure 4b, c).

**Figure 4.**
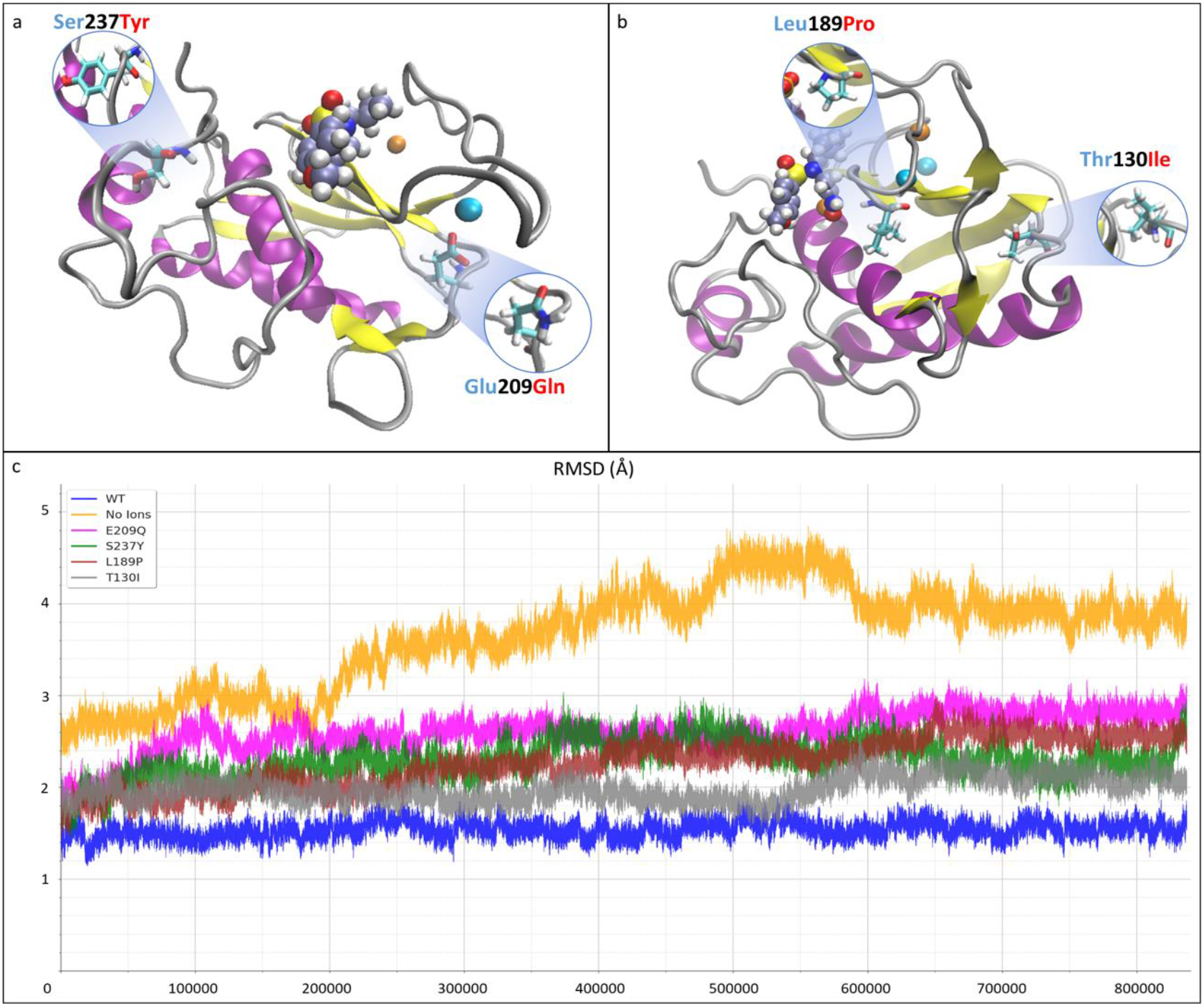
The tertiary structure of the catalytic domain of MMP20, based on the PDB:2JSD NMR model. a: The WT protein structure with the 2 novel variants, c.625G>C; p.(E209Q) of Families 4 – 7 and c.710C>A; p.(S237Y) of Families 8 – 9, of the active site are presented in the inlays. b: The WT protein structure with the inlays showing 2 of the known variants of the active site of MMP20, c.389C>T; p.(T130I) and c.566T>C; p.(L189P). c: The root mean square deviation (RMSD) of each modelled variant during molecular dynamics (MD) simulations of 900ns. An increase of RMSD value corresponds to a loss of stability, with the WT MMP20 structure being the most stable and the MMP20 structure in the absence of structural zinc and calcium ions being the least stable.

*In-silico* saturation mutagenesis of the MMP20 active site performed by Rhapsody shows that there are regions of the protein that are significantly more likely to cause a pathogenic effect when the residues located there are altered (Supp. Figure 5). These regions largely correlate with the sites of the known and novel variants and have an increased polyphen-2 score (Supp. Figure 5). The Rhapsody analysis was limited to the catalytic domain of MMP20, because it relies on the availability of a tertiary structure. The results of the SDM analysis are presented in Supp. Table 11, showing the changes of free energy, residue occlusion and solvent accessibility for the WT and each mutant respectively.

## Discussion

Exome sequencing in a cohort of non-syndromic AI families revealed biallelic *MMP20* variants in ten autosomal recessive families. MMP20 is a protease that plays an essential role during the secretory and early transition stages of amelogenesis. Defects in MMP20 cause AI in both humans and mice, due to a failure to process, degrade and remove proteins from the extracellular matrix scaffold upon which the developing dental enamel is formed. The hypomaturation phenotype observed in these families fits with this hypothesis, with the dental enamel being of normal volume but characterised by a loss of translucency, discolouration and premature enamel loss. Post eruptive changes probably determine the pattern of premature enamel failure. The phenotypes observed in the families presented are consistent with those described in previous reports of AI due to *MMP20* variants.

The variant identified in Families 1-3, (c.954-2A>T), has been reported previously (Gasse et al., 2017; J. W. Kim et al., 2005; Prasad et al., 2015; J. Timothy Wright et al., 2011), but the effect on splicing has not been determined experimentally. The prediction that the mutation most likely results in the loss of exon 7 from the *MMP20* transcript (J. W. Kim et al., 2005), therefore, remains speculative. If confirmed, this would break the reading frame and lead to an abnormal transcript which would be expected to be subject to nonsense mediated decay.

The missense variant identified in Families 4 – 7, p.(E209Q), is novel and replaces a negatively charged glutamic acid residue with neutral glutamine. E209 is a coordinating ligand for calcium ion binding and is therefore critical to the function of MMP20 (Andreini et al., 2004; Arendt et al., 2007; Yamakoshi et al., 2013). Replacement with glutamine decreases protein stability relative to WT in the MD simulations (see Supp. Table 8). Inspection of the WT atomistic structure shows that E209 has a pair of carboxylate oxygen atoms oriented towards the nearby Ca^2+^ ion. In E209 the average distance is less than 4 Å for both MD repeats (Supp. Table 10), However, the mutant Q209 only has one oxygen; in one of the two duplicate MD trajectories the interaction between this amide oxygen and Ca^2+^ remains strong (an inter-atomic distance of 2.5 Å) but in the other it is completely disrupted (inter-atomic distance of 6.9Å), allowing it to form non-native hydrogen bonds with other residues (see Supp. Table 9).

The novel missense variant identified in Families 8 and 9 replaces a small serine residue with a larger aromatic tyrosine, p.(S237Y) in a highly conserved part of the catalytic domain of MMP20. Correspondingly, the SDM analysis identifies an increase in the folding free energy of the protein relative to the WT (Supp. Table 11), implying the variant is less stable. Moreover, in the MD simulations the larger more hydrophobic Y237 results in an increased RMSD (2.35 Å) relative to the WT (1.55 Å). In these trajectories, the main backbone hydrogen bond formed by S237 (with M244) is also formed by Y237. However, the number of side-chain hydrogen bonds is reduced in Y237 compared to S237 (see Supp. Table 9), because the corresponding tyrosine oxygen protrudes too far into the solvent to participate in inter-atomic interactions within the protein.

In Family 10, the delins variant disrupts the intron 5 splice donor site and is therefore again likely to lead to a transcript that is subject to nonsense mediated decay. The novel c.1122A>C, p.(Q374H) variant changes a hydrophilic glutamine residue for a hydrophobic histidine in the C-terminal hemopexin-like domain, as shown in Figure 3a, disrupting the folding of the protein and potentially leading to a loss of function.

Microsatellites were used to determine whether the variants shared in each of the three family groups were founder alleles, implying these families are related, rather than that these sites are mutation hotspots. Family 1 is from Costa Rica, the population of which is largely of mixed European and Indigenous American descent. Families 2 and 3 are Caucasian European families from the UK. The c.954-2A>T variant has been reported previously in at least six families from France (Gasse et al., 2017; Prasad et al., 2015) and one from the USA (J. W. Kim et al., 2005). Ethnicity is given as Caucasian by Prassad and co-workers but is not given in the remaining reports. Our haplotype analysis shows that this variant is present on three different chromosomal backgrounds, but elements of the haplotype, in particular the genotype of D11S4108 100kb from *MMP20*, appear conserved between families. This suggests they may be related, but through a distant common ancestor. The picture is similar for Families 4, 5, 6 and 7, UK families of Pakistani origin. In contrast, Families 8 and 9 from Oman share an identical haplotype across the region tested. These findings therefore suggest that *MMP20* c.710C>A represents a relatively recent founder mutation in the Omani population, while c.954-2A>T and c.625G>C result from older founder mutations in the Caucasian European and Pakistani populations.

In addition to the novel p.(E209Q) and p.(S237Y) variants, we also performed MD for known variants p.(L189P) and p.(T130I) to see if the simulations could provide equivalent structural insight into loss of function. Other variants within the active site of MMP20 that change the ability of zinc ions to form co-ordination bonds were not modelled, because this will either lead to substantial structural distortion (as seen with MD in the absence of calcium or zinc ions), or destabilise the key zinc metal centre essential for catalysis.

For p.(L189P), while there is only a modest increase in RMSD relative to the WT, simulation of the behaviour of the bound ligand in the WT and variant shows that replacing bulky lysine with more compact proline changes the shape of the active site. This leads to substantial structural rearrangement of the ligand, which is likely to affect substrate recognition and catalytic activity of the protease against natural substrates.

The only pathogenic missense variant modelled that did not show a significantly larger RMSD than the WT was p.(T130I). While minor changes in hydrogen bonding interactions were detected (Supp. Table 9), no substantial changes in overall protein structure or perturbations to inter-atomic interactions with the ligand or metal ions were observed. Consequently, our MD simulations do not provide a structural mechanism for why this variant should be pathogenic. However, p.(T130I) is located at the edge of the catalytic domain, close to regions of the protein that are absent from the available structure and which therefore do not feature in these simulations.

The 14 *MMP20* variants previously reported to cause AI are shown in Figure 3, together with those newly described herein, indicating their location in the gene and protein. Previously reported pathogenic variants consisted of seven missense variants, two premature stop codons, two frameshifts and two putative splice variants. Here we report a further three missense and one splice variants, consistent with and extending the spectrum of mutations in *MMP20* causing autosomal recessive AI. The three splice variants have not been verified beyond *in-silico* prediction and therefore remain likely but unproven. These 17 variants, and in particular the missense variants, cluster primarily in or near to the zinc dependent peptidase domain (Figure 3). Our analyses suggest that pathogenic mutations in this domain alter protein stability, while others (Y. J. Kim et al., 2017) have shown that variants in the catalytic domain can lead to reduction or complete loss of enzymatic function. The combination of recessive inheritance, missense variants that reduce or abolish function and the lack of significant difference between phenotypes associated with missense and nonsense variants, all point to a loss of function phenotype, where lack of functional MMP20 gives rise to a consistent hypomaturation AI phenotype.

In summary we have identified ten families segregating AI due to four novel and one known mutations in *MMP20*, and reviewed previously reported mutations, raising the total number of AI causing *MMP20* variants to eighteen. This expands the spectrum of *MMP20* variants implicated in AI and confirms the association with a hypomaturation phenotype. Haplotype analysis suggests that the c.710C>A, p.(S237Y), c.625G>C, p.(E209Q) and c.954-2A>T, p.(I319Ffs*19) variants may be relatively common founder mutations in the Omani, Pakistani and Caucasian populations respectively. Biochemical modelling and molecular dynamic analyses show that missense mutations in the catalytic domain alter protein stability, suggesting that this form of AI is the result of loss of function of MMP20.

## Acknowledgements

The authors thank the families involved in this study. This work was supported by the Welcome Trust (grant 093113). GN was funded by a PhD scholarship funded by Leeds Dental School, Faculty of Medicine and Health, University of Leeds. CELS was funded by a Wellcome Trust Institutional Strategic Support Fund award. Funding for Open Access publication charges for this article was provided by the Wellcome Trust. This work was undertaken on ARC3, part of the High-Performance Computing facilities at the University of Leeds, UK.

