## Supplementary Materials for "Spectrum of pathogenic variants and multiple founder effects in amelogenesis imperfecta associated with *MMP20*"

Supplemental Data Contents:

**Supplementary Methods**

1. Detailed protocol for molecular dynamic simulations using AMBER

**Supplementary Tables**

1. **<u>Supplementary table 1.</u>** All known variants of *MMP20.* Data obtained from the AI - LOVD database
2. **<u>Supplementary table 2.</u>** Primers used for PCR and microsatellite markers used for genotyping.
3. **<u>Supplementary table 3.</u>** Coverage statistics for the WES of families 1 – 10.
4. **<u>Supplementary table 4.</u>** Family number, place of origin, number of affected members of each family recruited for this study. The coding and protein change, the genomic coordinates, CADD score and gnomAD frequency of each variant are also presented.
5. **<u>Supplementary table 5:</u>** MMP20 sequences used for conservation analysis. Databases last accessed 15 Dec 2019.
6. **<u>Supplementary table 6.</u>** Scores obtained from various the pathogenicity prediction software for the variants found in this study.
7. **<u>Supplementary table 7</u>:** Detailed results of the microsatellite analysis for families 1 – 9.
8. **<u>Supplementary table 8</u>**: RMSD average values for the last 300 ns of each MD simulation, along with the respective standard error.
9. **<u>Supplementary table 9</u>**: Hydrogen bonds formed during the molecular dynamic simulations. The bonds that are unique for only the WT, only the No-ions or only the mutant model are in bold.
10. **<u>Supplementary table 10</u>**: Average distance of the $O_2$ and the neighbouring $Ca^{2+}$ of the 209 residue in each of the WT and mutant models.
11. **<u>Supplementary table 11</u>**: SDM predictions

**Supplementary Figures**

1. **Supplementary Figure 1.** Evolutionary conservation for each novel missense variant.
2. **Supplementary Figure 2.** Human Splicing Finder analysis of the c.809_811+12delACGgtaagattattainsCCAG variant from Family 10.
3. **Supplementary Figure 3.** RMSD of the variants included in the MD analysis, showing both repeats
4. **Supplementary Figure 4.** Atomic fluctuation of the residues of the catalytic domain of MMP20 and the change of structure of the No-ions model over the 900 ns of MD simulations
5. **Supplementary Figure 5.** Rhapsody score for each possible amino acid change of the active site of MMP20.

**Supplementary Methods:**

1. Detailed protocol for molecular dynamic simulations using AMBER:

The MD simulations were set up using the xLeap module of AMBER18 (Case et al., 2018), using the ff14SB protein forcefield (Maier et al., 2015). The zinc ions were represented using a bonded model of the interactions between the metal centre and the four co-ordinated amino acid residues (Peters et al., 2010). All systems were solvated with a 10 Å rectangular box of TIP3P water molecules and neutralized with $Cl^-$ ions. Equilibration used an initial energy minimisation, followed by 80ps of restrained MD during which the system is heated to 300 K with gradual release of restraints. An unrestrained production run of 400ns was then performed for the WT, the protein in the absence of structural metal ions and each of the four selected variants. We also ran independent repeat simulations, in which a different set of random atomic velocities corresponding to a temperature of 300 K was assigned at the beginning of the production runs. Analysis of the trajectories, including calculation of the root mean square deviation (RMSD), per residue atomic fluctuations, distance between atoms and hydrogen bonds occupancies were performed with the CPPTRAJ module of AMBER18. Trajectories were visualized using the VMD software (Humphrey et al., 1996).

**Supplementary Tables**

**<u>Supplementary table 1.</u>** All known variants of *MMP20.* Data obtained from the AI - LOVD database, http://dna2.leeds.ac.uk/LOVD/genes/MMP20.

| Variant | Transcript change | Protein change | Genomic coordinates, GRCh37 | References |
|---|---|---|---|---|
| 1 | c.102G>A | p.(W34*) | g.102495949C>T | Papagerakis et al. 2008 |
| | | | | Chan et al 2011 |
| 2 | c.103A>C | p.(=) | g.102495948T>G | Prasad et al. 2016 |
| | | | | Kim et al. 2020 |
| 3 | c.126+6T>G | p.(=) | g.102495919A>C | Prasad et al. 2016 |
| 4 | c.323A>G | p.(Y108C) | g.102487594T>C | Gasse et al. 2017 |
| 5 | c.359delA | p.(N120Ifs*3) | g.102487558delT | Gasse et al. 2013 |
| 6 | c.389C>T | p.(T130I) | g.102482620G>A | Gasse et al. 2013 |
| | | | | Kim et al. 2017 |
| | | | | Gasse et al. 2017 |
| | | | | Kim et al. 2020 |
| 7 | c.540T>A | p.(Y180*) | g.102480745A>T | Kim et al. 2017 |
| 8 | c.566T>C | p.(L189P) | g.102480719A>G | Gasse et al. 2017 |
| 9 | c.611A>G | p.(H204R) | g.102480674T>C | Wang et al. 2013 |
| 10 | c.616G>A | p.(D206N) | g.102480669C>T | Kim et al. 2020 |
| 11 | c.625G>C | p.(E209Q) | g.102480660C>G | This study |
| 12 | c.678T>A | p.(H226Q) | g.102479801A>T | Ozdemir et al. 2005 |
| | | | | Wright et al. 2011 |
| | | | | Kim et al. 2017 |
| 13 | c.710C>A | p.(S237Y) | g.102479769G>T | This study |
| 14 | c.809_811+12 delACGgtaagattatta insCCAG | p.(?) | g.102479656_102479660+12 delTGCcattctaataatinsGGTC | This study |
| 15 | c.910G>A | p.(A304T) | g.102477309C>T | Lee et al. 2010 |
| | | | | Gasse et al. 2017 |
| 16 | c.954-2A>T | p.(I319Ffs*19) | g.102465490T>A | Kim et al. 2005 |
| | | | | Wright et al. 2011 |
| | | | | Prasad et al. 2016 |
| | | | | Gasse et al. 2017 |
| | | | | This study |
| 17 | c.1054G>A | p.(E352K) | g.102465388C>T | Seymen et al. 2015 |
| 18 | c.1122A>C | p.(Q374H) | g.102464295T>G | This study |

**Supplementary table 2.** Primers used for PCR and microsatellite markers used for genotyping.

| Target | Forward primer | Primer Tm (˚C) | Reverse primer | Primer Tm (˚C) | Expected product size (bp) |
|---|---|---|---|---|---|
| *MMP20* exon 4: c.625G>C | CTGTAATATGATGCGCCCCT | 57 | AGTTAAAGGGTGGCTTGGGA | 58 | 298 |
| *MMP20* exon 5 - intron 5: c.710C>A, also: c.809_811+12 delACGgtaagattatta insCCAG | GGGTTAACTGTAATGTGGGCAT | 57 | TGCACTTTCTTTAATTCGGAGA | 53 | 338 |
| *MMP20* exon 7: c.954-2A>T | GCAAGAGCAAAGGGCATTTA | 56 | ATGACTGGAAAAATGCTGGC | 56 | 350 |
| *MMP20* exon 8: c.1122A>C | CATTTAAGAGCCTTCATAGAAATCTT | 52 | TTCTTTCGTGGAAGGGTTTA | 53 | 326 |
| D11S940 | TCATCCCCAAATGCTCAG | 53 | GGAATCAAACTTCACATAGGAGG | 54 | ~183 |
| D11S1339 | ATGGCCTTGGAAAAATATC | 48 | GGGTGTAACCAGTTCTTCAG | 54 | ~122 |
| D11S4108 | TGGCAAGTGGCAGGAT | 55 | GCCCATAGATGGATGAGTAGA | 54 | ~113 |
| D11S4159 | CCGGAGAGCAGTTTGTGT | 56 | ATTCGGAGCCACTCCCT | 57 | ~180 |
| D11S4161 | ACTCCCTGAAGTGTGCTAGGAAG | 60 | GTTTTAGGTTTCCATCTGGCAA | 54 | ~126 |
| D11S4951 | ATGGGTATACACCCAGCAAA | 55 | AACTGTGATTTTAAAAGATAATGCC | 50 | ~130 |

**Supplementary table 3.** Coverage statistics for the WES of families 1 – 10.

| Family | Mean Coverage | % Bases covered >4 | % Bases covered >14 | % Bases covered >19 |
|---|---|---|---|---|
| 1 | 54.07 | 93.6 | 84.9 | 80.1 |
| 2 | 89.56 | 99.4 | 97.4 | 95.8 |
| 3 | 55.34 | 98.6 | 90.0 | 83.8 |
| 4 | 60.01 | 93.6 | 86.7 | 83.0 |
| 5 | 94.43 | 95.2 | 90.2 | 88.0 |
| 6 | 121.00 | >99.3 | 99.3 | 98.1 |
| 7 | 51.64 | 94.3 | 86.0 | 81.1 |
| 8 | 58.10 | 93.9 | 85.8 | 81.4 |
| 9 | 47.41 | 92.8 | 84.2 | 79.3 |
| 10 | 94.38 | 91.6 | 86.8 | 84.5 |

**Supplementary table 4.** Family number, place of origin, number of affected members of each family recruited for this study. The coding and protein change, the genomic coordinates, CADD score and gnomAD frequency of each variant are also presented.

| Family | Origin | Affected | Transcript change | Protein change | Genomic Coordinates, GRCh37 | CADD | gnomAD |
|---|---|---|---|---|---|---|---|
| 1 | Costa Rica | 1 | c.954-2A>T | p.(I319Ffs*19) | g.102465490T>A | 24.5 | rs140213840 - 0.001082 |
| 2 | UK – Caucasian | 1 | | | | | |
| 3 | UK – Caucasian | 3 | | | | | |
| 4 | UK – Pakistani | 1 | c.625G>C | p.(E209Q) | g.102480660C>G | 27.2 | rs199788797 - 0.00005685 |
| 5 | UK – Pakistani | 1 | | | | | |
| 6 | UK – Pakistani | 3 | | | | | |
| 7 | UK – Pakistani | 2 | | | | | |
| 8 | Oman | 2 | c.710C>A | p.(S237Y) | g.102479769G>T | 28.4 | Absent |
| 9 | Oman | 1 | | | | | |
| 10 | UK – Caucasian | 1 | c.809_811+12 delACGgtaagattatta insCCAG | p.(?) | g.102479656_102479670delinsCTGG | N/A | Absent |
| | | | c.1122A>C | p.(Q374H) | g.102464295T>G | 18.81 | Absent |

**<u>Supplementary table 5:</u>** MMP20 sequences used for conservation analysis. Databases last accessed 15 Dec 2019.

| Species | Accession No | Amino acid sequence |
|---|---|---|
| *Homo sapiens* (Human) | ENST00000260228 | MKVLPASGLAVFLIMALKFSTAAPSLVAASPRTWRNNYRLAQAYLD<br>KYYTNKEGHQIGEMVARGSNSMIRKIKELQAFFGLQVTGKLDQTT<br>MNVIKKPRCGVPDVANYRLFPGEPKWKKNTLTYRISKYTPSMSSVE<br>VDKAVEMALQAWSSAVPLSFVRINSGEADIMISFENGDHGDSYPFD<br>GPRGTLAHAFAPGEGLGGDTHFDNAEKWTMGTNGFNLFTVAAHE<br>FGHALGLAHSTDPSALMYPTYKYKNPYGFHLPKDDVKGIQALYGPR<br>KVFLGKPTLPHAPHHKPSIPDLCDSSSSFDAVTMLGKELLLFKDRIFW<br>RRQVHLRTGIRPSTITSSFPQLMSNVDAAYEVAERGTAYFFKGPHY<br>WITRGFQMQGPPRTIYDFGFPRHVQQIDAAVYLREPQKTLFFVGDE<br>YYSYDERKRKMEKDYPKNTEEEFSGVNGQIDAAVELNGYIYFFSGPK<br>TYKYDTEKEDVVSVVKSSSWIGC |
| *Pan troglodytes* (Chimpanzee) | ENSPTRT00000007863 | MKVLPASGLAVFLIMALKFSTAAPSLVAASPRTWRNNYRLAQAYLD<br>KYYTNKEGHQIGEMVARGSNSMIRKIKELQAFFGLQVTGKLDQTT<br>MNVIKKPRCGVPDVANYRLFPGEPKWKKNILTYRISKYTPSMSSVEV<br>DKAVEMALQAWSSAVPLSFVRINSGEADIMISFENGDHGDSYPFD<br>GPRGTLAHAFAPGEGLGGDTHFDNAEKWTMGTNGFNLFTVAAHE<br>FGHALGLAHSTDPSALMYPTYKYKNPYGFHLPKDDVKGIQALYGPR<br>KAFLGKPTLPHAPHHKPSIPDLCDSSSSFDAVTMLGKELLLFKDRIFW<br>RRQVHLRTGIRPSTITSSFPQLMSNVDAAYEVAERGTAYFFKGPHY<br>WITRGFQMQGPPRTIYDFGFPRHVQQIDAAVYLREPQKTLFFVGDE<br>YYSYDERKKKMGKDYPKNTEEEFSGVNGQIDAAVELNGYIYFFSGPK<br>TYKYDTEKEDVVSVVKSSSWIGC |
| *Otolemur garnettii* (Bushbaby) | ENSOGAT00000015251 | MKMLPASGLAVLLITALKLSTAAPSLFTATPRTWRNNYHLAQEYLDK<br>YYTKKGGHQIGEMVARGGNSMVKKIKELQAFFGLQVTGKLDATTM<br>DVIKRPRCGVPDVANYRLFPGEPKWKKNTLTYRISKYTPSMSSAEVD<br>TAIEMALRAWSSAVPLNFVRVNTGEADIMISFETGDHGDSYPFDGP<br>RGTLAHAFAPGEGLGGDTHFDNAEKWTLGMNGFNLFTVAAHEFG<br>HALGLAHSTDPSALMYPTYKYQNPYGFHLPMDDVKGIQALYGPRK<br>PFLGKPTMPHGPPHNPPTPDLCDSSSSFDAVTMLGKELLFFKDRIF<br>WRRQVHLPTGIRPSTITSSFPQLMSNVDAAYEVAERGTAYFFKGPH<br>YWITRGFQMQGPPRTIYDFGFPRHVQRIDAAVYLREPRKTLFFVGD<br>EYYSYDERKRKMEKDYPKNTEEEFSGVNGQIDAAVELNGYIYFFSGP<br>KAYKYDMEKEDVVSVVKSSSWVGC |
| *Mus musculus* (Mouse) | ENSMUST00000034487 | MKVLPASGLAVLVTALKFATADPNLLAATPRTFRSNYHLAQAYLDKY<br>YTKKGGPQAGEMVARESNPMIRRIKELQIFFGLKVTGKLDQNTMN<br>VIKKPRCGVPDVANYRLFPGEPKWKKNILTYRISKYTPSMSPTEVDK<br>AIQMALHAWSTAVPLNFVRINSGEADIMISFETGDHGDSYPFDGPR<br>GTLAHAFAPGEGLGGDTHFDNAEKWTMGTNGFNLFTVAAHEFGH<br>ALGLGHSTDPSALMYPTYKYQNPYRFHLPKDDVKGIQALYGPRKIFP<br>GKPTMPHIPPHKPSIPDLCDSSSSFDAVTMLGKELLFFKDRIFWRRQ<br>VHLPTGIRPSTITSSFPQLMSNVDAAYEVAERGIAFFFKGPHYWVTR<br>GFHMQGPPRTIYDFGFPRHVQRIDAAVYLKEPQKTLFFVGEEYYSYD<br>ERKKKMEKDYPKNTEEEFSGVSGHIDAAVELNGYIYFFSGRKTFKYD<br>TEKEDVVSVVKSSSWIGC |
| *Ictidomys tridecemlineatus* (Squirrel) | ENSSTOT00000027785 | MKVLPASGLAVLLITALKFSTAAPSLFAATPRTWRNNYHLAQEYLDK<br>YYTKKGGHPVGEMAARGGNAMVKKIKELQAFFGLQVNGKLDQNT<br>MDVIKRPRCGVPDVANYRLFPGEPKWKKNTLTYRIAKYTSSMRPIEV<br>EKAVEMALQAWSSAVPLSFVRINSGEADIMISFETGDHGDSYPFDG<br>PRGTLAHAFAPGEGLGGDTHFDNAEKWTMGMNGFNLFTVAAHE |

| | | |
|---|---|---|
| | | FGHALGLAHSSDPTALMYPTYKYQNPYGFRLPKDDVKGIQALYGPR<br>KPFLGKPTIPHVPPHKPSNPDPCDSRASFDAVTMLGKELLFFRDRIF<br>WRRQVHVPAGIRPSTITSSFPQLMSNVDAAYEVAERGTAYFFKGPH<br>YWITRGFQMQGPPRTIYDFGFPRHVQRIDAAVYLKKPQKTLFFVGD<br>EYYSYDERKRKMDKDYPKNTEEEFSGVSGQIDAAVELNGHIYFFSGP<br>KTYKYDTEKEDVVSVVKSSSWIGC |
| *Canis lupus familiaris* (Dog) | ENSCAFT00000023926 | MTVLPMCGLALLLGAALEFCTAAPSVSAAAPRTTQNKYHLAQAYLD<br>KYYTSKAGPQVGEMGAPGGRALIKKIKELQAFFGLRITGKLDRPTMD<br>MIKRPRCGVPDVANYRLFPGEPKWKKNTLTYRISKYTSSMSPAEVD<br>KAVEMALQAWGSAVPLSFIRVNSGEADIMISFETGDHGDSYPFDGP<br>RGTLAHAFAPGEGLGGDTHFDNAEKWTMGMNGFNLFTVAAHEF<br>GHALGLAHSTDPSALMYPTYKYQHPYGFHLPKDDVKGIQALYGPRK<br>TLLGKPTVPHAPPQSPSIPDLCDSSSSFDAVTMLGKELLLFRDRIFWR<br>RQVHLMAGIRPSTITSSFPQLMSNVDAAYEVAERGTAYFFKGPHYW<br>ITRGFQMQGPPRTIYDFGFPRYVQRIDAAVYLKDVQKTLFFVGDEYY<br>SYDERKRKMEKDYPKNTEEEFSGVNGQIDAAVELNGYIYFFSGPKAY<br>KYDTEKEDVVSVLKSSSWIGC |
| *Erinaceus europaeus* (Hedgehog) | XM_007520728 | MKVLPTSGFAVLLIMALKLSTAAPSLFAATPRTSRNNYHLAQEYLDR<br>YYTKKEEYQIGEMVARGHNSMIKKIKELQAFFGLQITGKLDRTTMDV<br>IKKPRCGVPDVANYRLFPGEPKWKKNTLTYRISKYTSSMTSAEVDKA<br>VEMALQAWSSAVPLNFVKINSGEADIMISFETGDHGDSYPFDGPRG<br>TLAHAFAPGEGLGGDTHFDNAEKWTMGMNGFNLFTVAAHEFGH<br>ALGLAHSTDPSALMYPTYKYQHPYGFHLPKDDVKGIQALYGPRKTFP<br>GKPTIPFSPPHNPSIPDLCDTSSSFDAVTMLGKELLFFRDRIFWRRQV<br>HLPGGIRPSTITSSFPQLMSNVDAAYEVAERGTAYFFKGPHYWITRG<br>FRMQGPPRTIYDFGFPRYVQRIDAAVYLKDAQKTLFFVGDEYYSYDE<br>RKGKMEKDYPKNTEEEFSGVNGQIDAAVELNGYIYFFSGPKAYKYDT<br>EKEDVVSVVKSSSWIGC |
| *Loxodonta africana* (Elephant) | ENSLAFT00000010998 | MKVLPASGLAVLFITTLKFSTAAPSLFAATSRTSRNNYQLAQAYLDKY<br>YTKEGGHQIGEMVARGGNAMVKKIKELQAFFGLKVTGKLDQLTIDV<br>IKKPRCGVPDVANYRLFPGEPKWKKNTLTYRISKYTSSMSSADVDKA<br>IEMALQAWSSAIPLSFVKLNTGEADIMISFETGDHGDSYPFDGPRGT<br>LAHAFAPGEGLGGDTHFDNAEKWTMGMNGFNLFTVAAHEFGHA<br>LGLAHSTDPSALMYPTYKYQHPYGFRLPKDDVKGIQALYGPRKTFPG<br>KPTVPHGPPQNPSTPDLCDSSSSFDAVTMLGKELLFFKDRIFWRRQ<br>AHLMAGIRPSTITSSFPQIMANVDAAYEVAERGAAYFFKGPHYWVT<br>RGFQMQGPPRSIYDFGFPRFVQQIDAAVYLKNAQKTLFFVGDEYYS<br>YDERKGKMEKDYPKNTEEEFSGVSGQIDAAVELNGYIYFFSGPKAYK<br>YDTEKEDVVSVLKSSSWIGC |
| *Xenopus tropicalis* (Frog) | ENSXETT00000064032 | YLDKYYSDRGTMRVAEMVADDVSMSRKIRKMQKFFGLQVTGKLD<br>HSTLAVMQKPRCGMPDLANYHVFPGEPKWQRSSLTYRITKYTSSLS<br>TQDVDRAVDLGLKAWSDAAPLNFIKTTQGEADIMISFESGDHGDSY<br>PFDGPRGTLAHAFAPGEGLGGDTHFDNAERWTTGKNGFNLFTVAA<br>HEFGHALGLGHSSDPSALMYPTYRYQHPIGFQLPTDDVKGIQALYG<br>TKGIGKEKPMAPQQPANKPDQCDPNLSFDAVTVLGNELLFFKLTRS<br>FWRRQAPLTNIGPSPIASSFPQLMSNVDAAYEVAEQGTAYFFKGLIG<br>PHYWATRGLQMQGHPRTIYDFGFPRHVQKIDAAVHLKNSRKTLFF<br>VGDDYYSYDETKREMEDDYPKSIDDEFTGVEGNIDAAVEVNGFIYFF<br>SGPKAYKYDTEKEDVVNIVKSSSWIGC |

**Supplementary table 6.** Scores obtained from various the pathogenicity prediction software for the variants found in this study.

| Variant | PROVEAN | SIFT | CADD v1.3 | MutPred2 |
|---|---|---|---|---|
| c.625G>C, p.(E209Q) | Deleterious (-2.84) | Damaging (0.002) | 27.2 | 0.84 |
| c.710C>A, p.(S237Y) | Deleterious (-5.83) | Damaging (0) | 28.4 | 0.818 |
| c.809_811+12 delACGgtaagattatta insCCAG, p.(?) | N/A | N/A | N/A | N/A |
| c.954-2A>T, p.(I319Ffs*19) | N/A | N/A | 24.5 | N/A |
| c.1122A>C, p.(Q374H) | Neutral (-1.38) | Damaging (0.016) | 18.81 | 0.268 |

**Supplementary table 7:** Detailed results of the microsatellite analysis.

| | | D11S940 | | D11S1339 | | D11S4108 | | D11S4161 | | D11S4159 | | |
|---|---|---|---|---|---|---|---|---|---|---|---|---|
| Family No | Family member | Allele A | Allele B | Allele A | Allele B | Allele A | Allele B | Allele A | Allele B | Allele A | Allele B | Affected |
| 1 | II:1 | 175 | 175 | 129 | 139 | 109 | 109 | 120 | 126 | 171 | 171 | YES |
| 2 | II:1 | 162 | 175 | 113 | 119 | 109 | 109 | 126 | 126 | 171 | 171 | NO |
| | II:2 | 162 | 179 | 113 | 139 | 109 | 109 | 126 | 126 | 171 | 171 | NO |
| | III:1 | 162 | 162 | 113 | 113 | 109 | 109 | 126 | 126 | 171 | 171 | YES |
| | III:2 | 175 | 179 | 119 | 139 | 109 | 109 | 126 | 126 | 171 | 171 | NO |
| 3 | II:5 | 162 | 181 | 113 | 129 | 109 | 109 | 126 | 126 | 171 | 171 | NO |
| | III:1 | 175 | 181 | 113 | 129 | 109 | 121 | 126 | 138 | 171 | 171 | NO |
| | III:2 | 162 | 175 | 113 | 129 | 109 | 109 | 126 | 126 | 171 | 171 | YES |
| | III:3 | 162 | 175 | 113 | 129 | 109 | 109 | 126 | 126 | 171 | 171 | YES |
| | III:4 | 181 | 181 | 129 | 129 | 109 | 109 | 126 | 126 | 171 | 179 | NO |
| 4 | IV:1 | 181 | 181 | 133 | 133 | 121 | 121 | 126 | 126 | 171 | 171 | YES |
| | IV:2 | 181 | 181 | 133 | 133 | 121 | 121 | 126 | 126 | 171 | 171 | YES |
| 5 | V:1 | 181 | 181 | 133 | 133 | 121 | 121 | 126 | 126 | 171 | 171 | YES |
| 6 | II:1 | 183 | 183 | 146 | 146 | 109 | 109 | 126 | 126 | 179 | 179 | YES |
| 7 | IV:2 | 175 | 183 | 113 | 146 | 109 | 109 | 138 | 138 | 179 | 179 | YES |
| | IV:4 | 183 | 183 | 146 | 146 | 109 | 109 | 138 | 138 | 179 | 179 | YES |
| 8 | III:2 | 175 | 183 | 129 | 139 | 109 | 109 | 126 | 126 | 171 | 171 | NO |
| | IV:1 | 175 | 175 | 129 | 129 | 109 | 109 | 126 | 126 | 171 | 171 | YES |
| 9 | III:1 | 175 | 183 | 139 | 146 | 109 | 121 | 126 | 138 | 171 | 179 | NO |
| | III:2 | 170 | 183 | 129 | 146 | 109 | 121 | 126 | 126 | 171 | 179 | NO |
| | IV:1 | 183 | 183 | 129 | 129 | 109 | 121 | 126 | 138 | 171 | 179 | NO |
| | IV:2 | 183 | 183 | 129 | 146 | 109 | 121 | 126 | 138 | 171 | 179 | NO |
| | IV:3 | 183 | 183 | 146 | 146 | 109 | 109 | 138 | 138 | 179 | 179 | YES |
| | IV:4 | 175 | 183 | 129 | 139 | 111 | 121 | 126 | 126 | 171 | 171 | NO |
| | IV:5 | 183 | 183 | 146 | 146 | 109 | 109 | 138 | 138 | 179 | 179 | YES |

**Supplementary table 8**: RMSD average values for the last 300 ns of each MD simulation, along with the respective standard deviation, for each repeat and their combination.

| | Average | Standard deviation |
|---|---|---|
| WT-1 | 1.74 | 0.11 |
| WT-2 | 1.36 | 0.14 |
| **WT** | **1.55** | **0.22** |
| No-ions-1 | 4.20 | 0.19 |
| No-ions-2 | 3.62 | 0.15 |
| **No-ions** | **3.91** | **0.34** |
| E209Q-1 | 2.43 | 0.10 |
| E209Q-2 | 3.17 | 0.19 |
| **E209Q** | **2.80** | **0.40** |
| S237Y-1 | 2.32 | 0.27 |
| S237Y-2 | 2.38 | 0.26 |
| **S237Y** | **2.35** | **0.27** |
| T130I-1 | 1.68 | 0.12 |
| T130I-2 | 2.53 | 0.14 |
| **T130I** | **2.10** | **0.45** |
| L189P-1 | 2.15 | 0.13 |
| L189P-2 | 2.92 | 0.17 |
| **L189P** | **2.53** | **0.41** |

**Supplementary table 9**: Key hydrogen bonds formed during the molecular dynamics simulations, expressed as a percentage occupancy over the 900ns MD trajectory. Empty cells indicate no occupancy detected. Grey cells indicate where either the donor or acceptor are not present in the sequence.

| | | E209Q | | WT | |
|---|---|---|---|---|---|
| Acceptor | Donor | 1st run | 2nd run | 1st run | 2nd run |
| Q209/E209 | W211 | 0.097 | 0.153 | 0.279 | 0.190 |
| Q209/E209 | S244 | 0.059 | 0.151 | 0.191 | 0.195 |
| D206 | Q209/E209 | 0.019 | 0.457 | 0.019 | 0.015 |
| P185 | Q209 | 0.586 | 0.050 | | |
| T188 | Q209 | 0.031 | | | |
| N207 | Q209 | 0.012 | 0.029 | | |
| Q209 | K210 | 0.003 | 0.041 | | |
| | | S237Y | | WT | |
| Acceptor | Donor | 1st run | 2nd run | 1st run | 2nd run |
| M244 | Y237/S237 | 0.344 | 0.336 | 0.431 | 0.544 |

| | | | | | |
|---|---|---|---|---|---|
| D262 | Y237/S237 | 0.070 | 0.125 | 0.164 | |
| Y237/S237 | D239 | 0.051 | 0.062 | 0.162 | 0.081 |
| D261 | Y237 | 0.026 | 0.054 | | |
| S237 | T238 | | | 0.093 | |
| D239 | S237 | | | 0.085 | 0.040 |
| S237 | H236 | | | 0.014 | |
| | | T130I | | WT | |
| Acceptor | Donor | 1st run | 2nd run | 1st run | 2nd run |
| I130 | M133 | 0.043 | 0.015 | | |
| N207 | T130 | | | 0.683 | 0.633 |
| T130 | S244 | | | 0.245 | 0.433 |
| | | L189P | | WT | |
| Acceptor | Donor | 1st run | 2nd run | 1st run | 2nd run |
| P189/L189 | D206 | 0.648 | 0.732 | 0.792 | 0.718 |

**Supplementary table 10**: Average distance and standard deviation of the $O_2$ and the neighbouring $Ca^{2+}$ of the 209 residue in each of the WT-E209 and Q209 models.

| Model | Average | Standard deviation |
|---|---|---|
| Q209:OE1-CA161_1st_run | 2.49 | 0.37 |
| Q209:OE1-CA161_2nd_run | 6.88 | 1.42 |
| WT:OE1-CA161_1st_run | 3.92 | 0.65 |
| WT:OE1-CA161_2nd_run | 2.82 | 0.77 |
| WT:OE2-CA161_1st_run | 2.50 | 0.37 |
| WT:OE2-CA161_2nd_run | 2.92 | 0.84 |

**Supplementary table 11**: SDM predictions of the changes in residue solvent accesibility (RSA%), residue occluded surface packing (OSP%), residue depth in the structure in A and free energy (ΔΔG).

| | E209Q | S237Y | T130I | L189P |
|---|---|---|---|---|
| WT_RSA(%) | 5.9 | 41.2 | 0.1 | 7.9 |
| MT_RSA(%) | 3.3 | 31.6 | 0.8 | 22.8 |
| **MT-WT_RSA(%)** | **-2.6** | **-9.6** | **0.7** | **14.9** |
| WT_DEPTH (Å) | 5.8 | 3.9 | 6.4 | 4.9 |
| MT_DEPTH (Å) | 6 | 4.3 | 6 | 4.6 |
| **MT-WT_Depth (Å)** | **0.2** | **0.4** | **-0.4** | **-0.3** |
| WT_OSP(%) | 0.45 | 0.28 | 0.47 | 0.48 |
| MT_OSP(%) | 0.49 | 0.28 | 0.53 | 0.35 |
| **MT-WT_OSP(%)** | **0.04** | **0** | **0.06** | **-0.13** |
| **Predicted ΔΔG** | **-0.15** | **0.36** | **0.7** | **-1.29** |

### Supplementary Figures

**Supplementary Figure 1.** Evolutionary conservation for each novel missense variant.

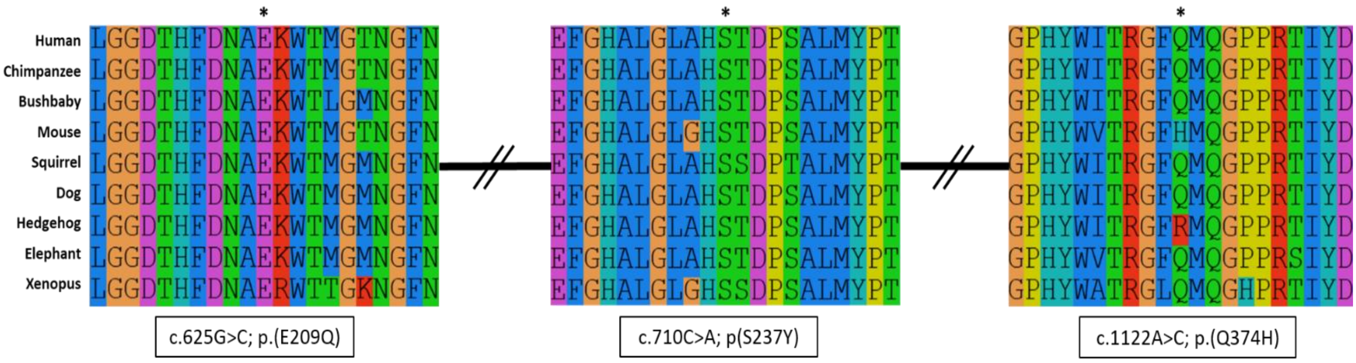


**Supplementary Figure 2.** Results of the Human Splicing Finder analysis of the c.809_811+12delACGgtaagattattainsCCAG variant from Family 10.

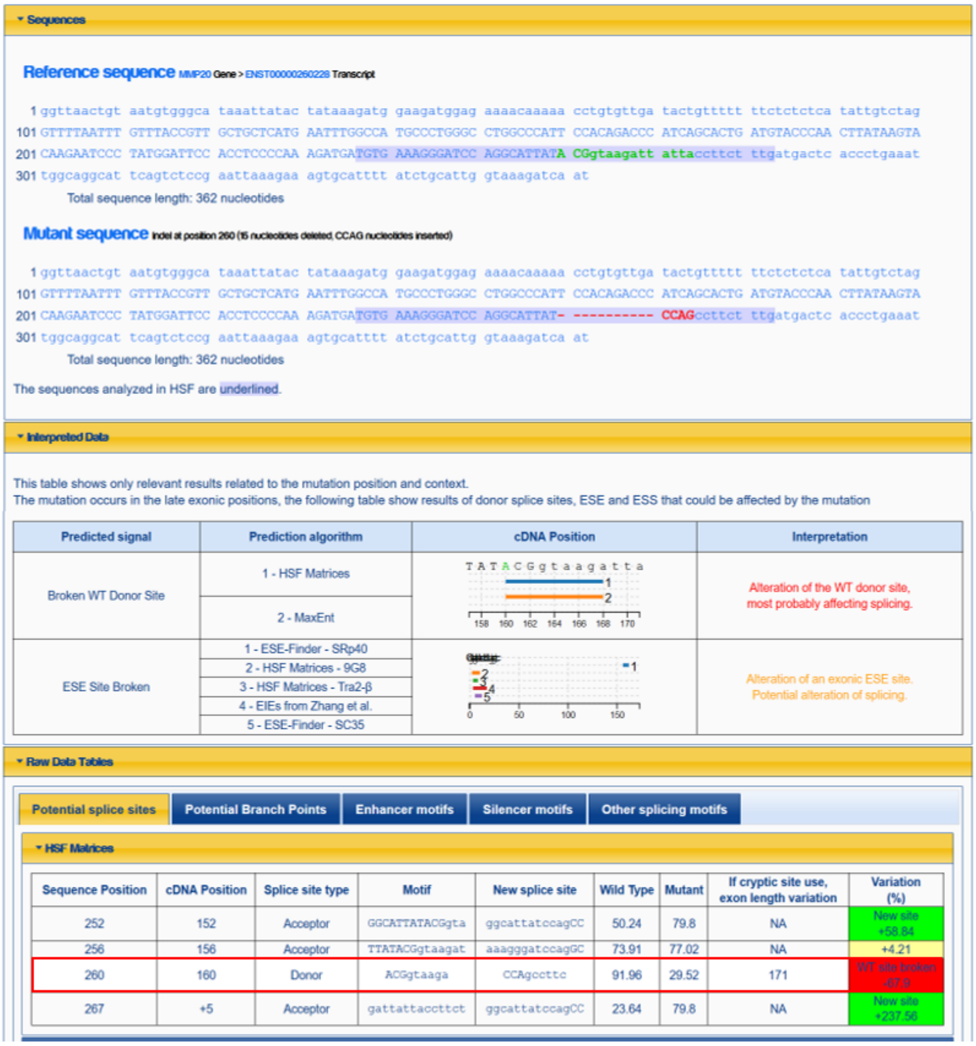

**<u>Supplementary Figure 3.</u>** RMSD of the variants included in the molecular dynamics analysis, showing both repeats. a: the novel variants described in this study b: the known variants from the literature. The WT and No-ions models are shown in both figures as a baseline for comparison.

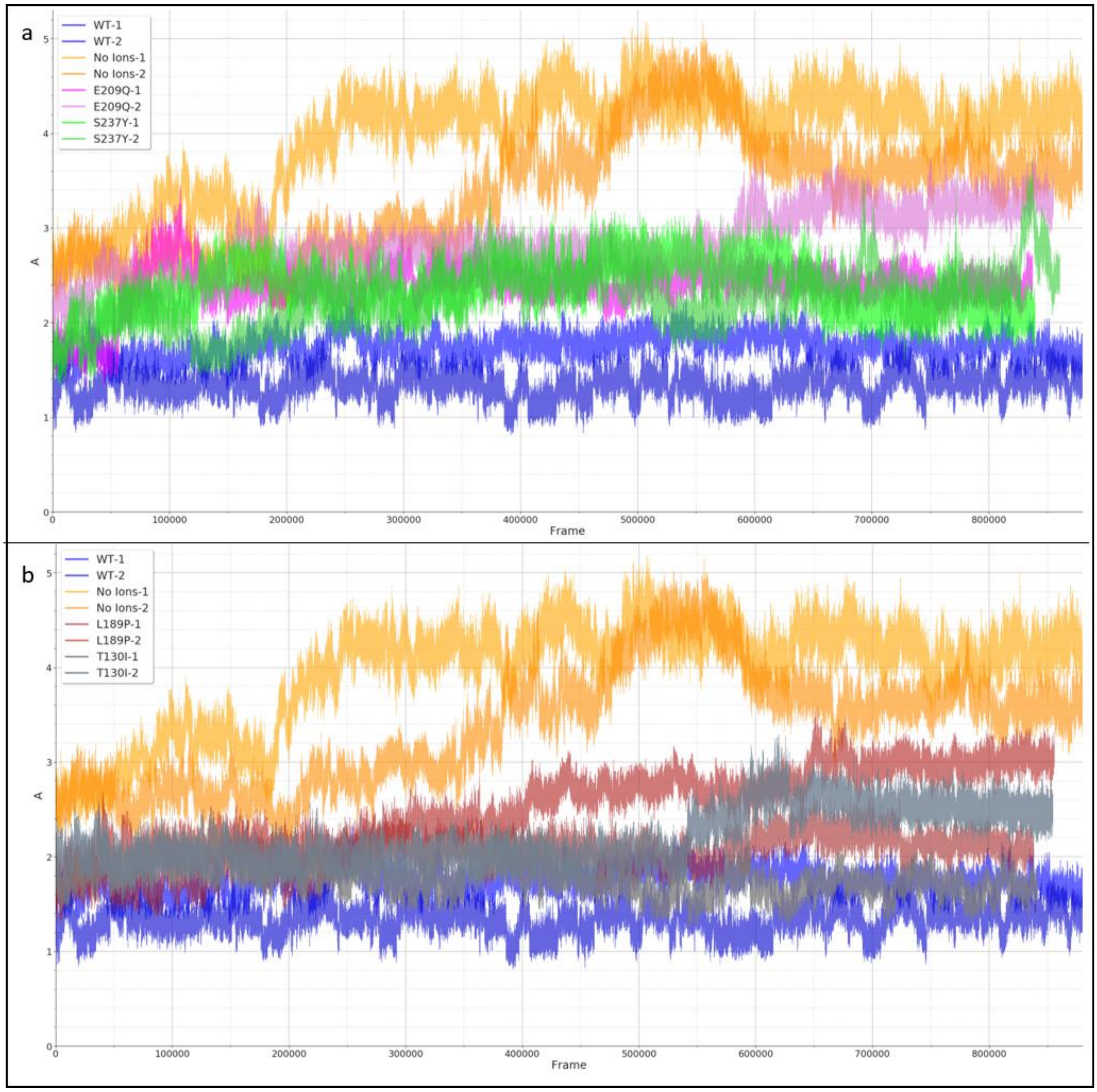

**Supplementary Figure 4.** a: Atomic fluctuation of the residues of the catalytic domain of MMP20. b, c: The structure of the No-ions model after the 900 ns of molecular dynamic simulations, the surface view (b) and the internal structure (c) at the start (red), midpoint (green) and end (blue) of the simulation.

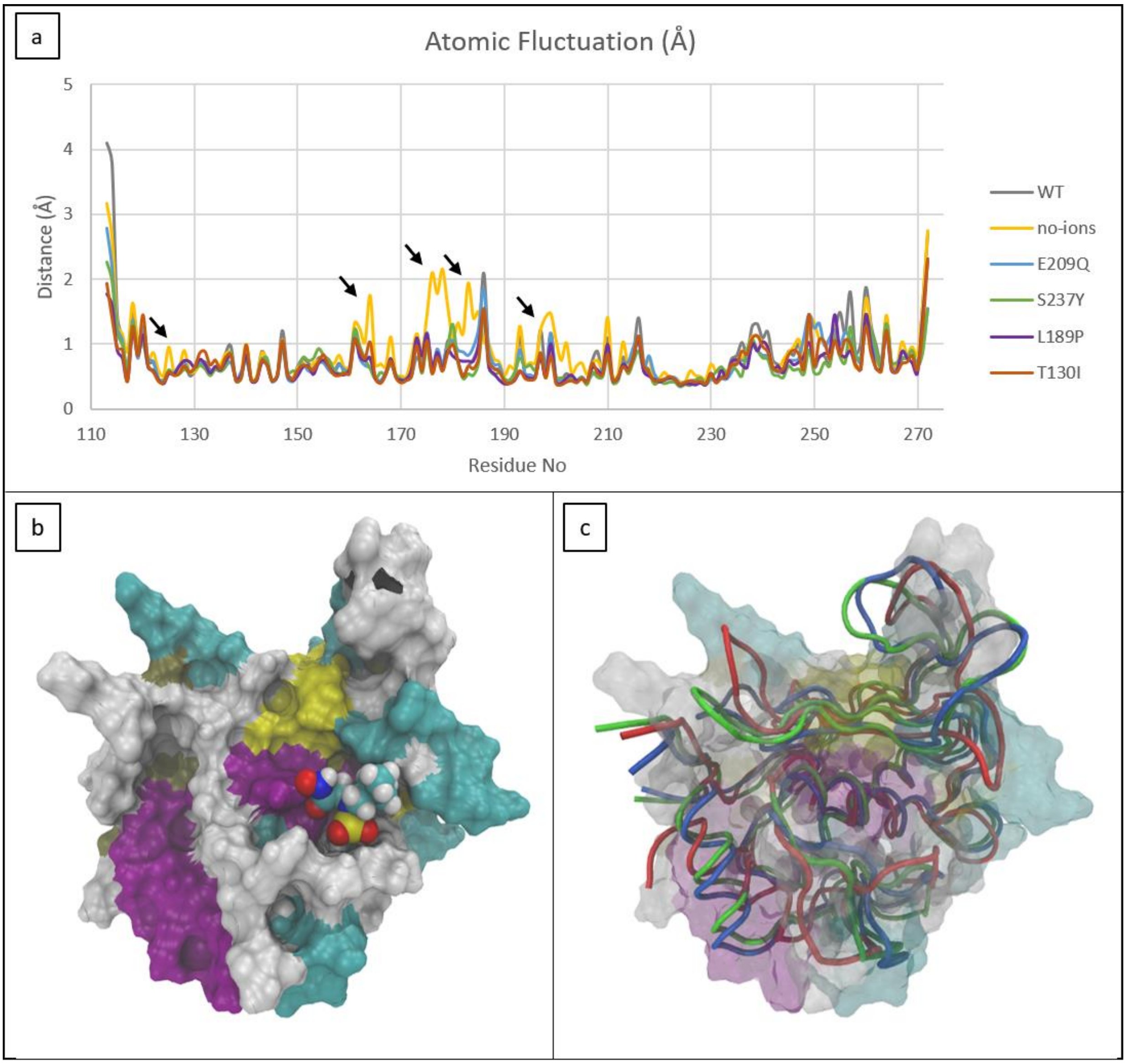

**<u>Supplementary Figure 5.</u>** Rhapsody score for each possible amino acid change of the active site of MMP20. Missense variants discussed in this study are indicated with a black arrow.

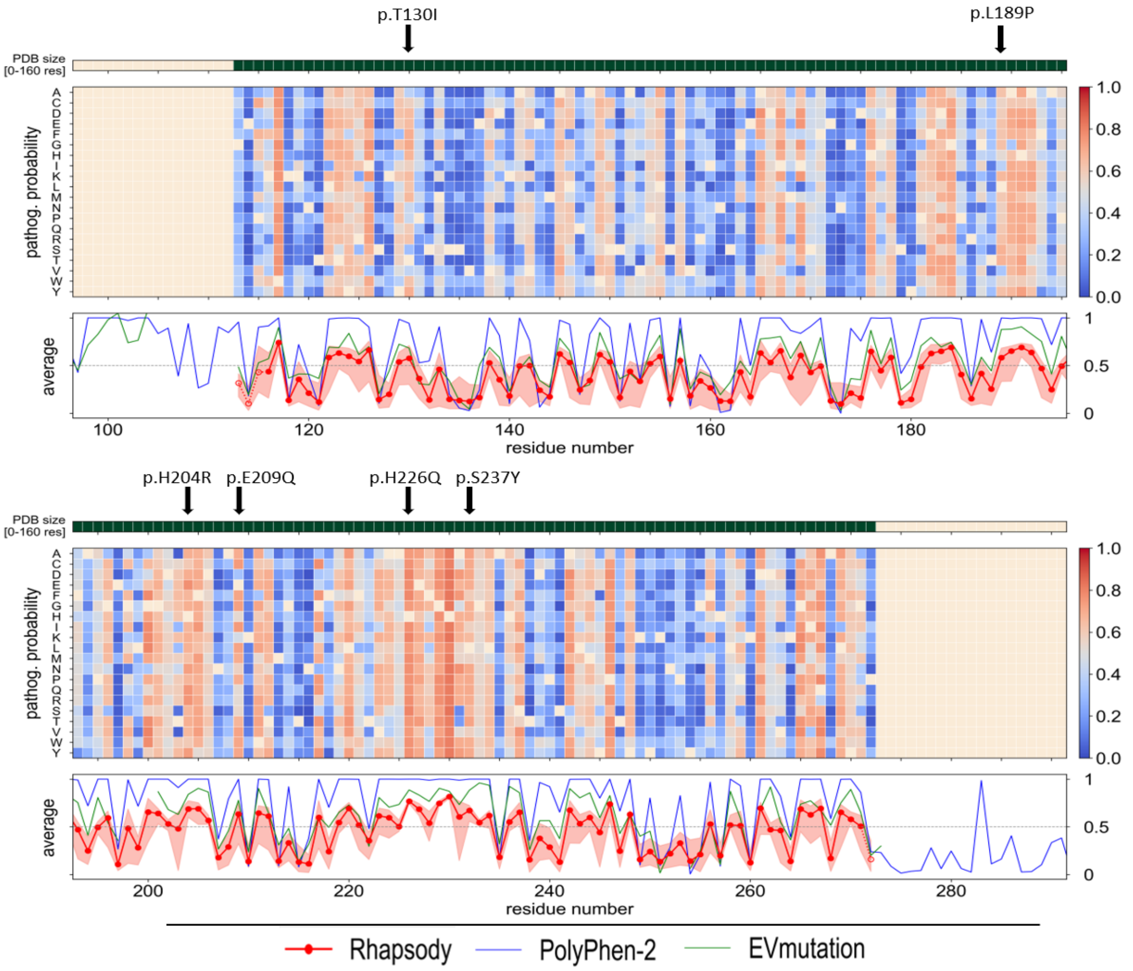
